# A massively parallel algorithm for finding non-existing sequences in genomes

**DOI:** 10.1101/709949

**Authors:** Marco Falda

## Abstract

We discuss a method for producing a set of absent words in a reference genome with a guaranteed Hamming distance along all positions and additional information about the number of mismatches, their location and the position of the best match. We implemented it exploiting the massively parallelism of modern GPUs hardware: the code is available at https://bitbucket.org/mfalda/cuda_keeseek/.

## I. INTRODUCTION

Non-existent sequences in genomes, also known as nullomers, have been considered for a number of different biomedical applications, for example they are thought to impact population genetics and they can be used as molecular tags or as specific adaptors for PCR. However, to the best of our knowledge, all algorithms proposed so far for nullomer generation are only focused on the detection of absent words in genomes, without providing any information about their distance in terms of number of mismatches, and they focus on words with a limited length. When such words are to be employed as barcodes or PCR primers, absent words must be distant enough to any position of the reference genome, and must possess "primer-like" features that allow them to be applied as primers.

In this paper we propose an algorithm for producing, for a given reference genome, a set of sufficiently long absent words in that genome (>= 18) with a guaranteed Hamming distance along all positions of the reference and additional information about the number of mismatches, their location and the position of the best match in the reference genome. The problem is not easy: The solution space provided by an arbitrary genome cannot be characterized in terms of a fitness function, therefore it is nearly impossible to design in a correct way an informed search exploiting heuristic properties. More formally, this is an 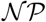-Complete problem, and this fact can be proven by reducing it from the Hamming Center Problem (HCP) [1]. HCP has been studied extensively in Theoretical Computer Science and also in Computational Biology, however most studies are focused on the related clustering problem, while the aim of this paper is to provide reasonable primers with fair biological properties. Meta-heuristics and parallel implementations with good practical running times have also been developed; the drawback of these approaches is that they cannot guarantee that an exact solution will be found.

Since the aim is quite practical, in that we just need to produce a number of potential good primers, we can exploit such requirements and move their selection before their exhaustive search in the genome; this means that we can apply Biology-driven criteria to our candidates in order to reduce their number *a priori*. The state-of-the-art parallel devices are now the Graphical Processing Units (GPUs). GPGPU computing is the use of a GPU together with a CPU to accelerate general-purpose scientific and engineering applications. It offers unprecedented application performance by offloading compute-intensive portions of the application to the GPU, while the remainder of the code still runs on the CPU. Preliminary tests on a optimized version exploiting the complex memory hierarchy of modern devices seem promising even on a consumer class device with a limited amount of resources with respect to the multi-threaded algorithm.

The article is organized as follows. Section II starts with a discussion of the temporal complexity of the problem at hand and it contains a proof of its 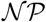-completeness. In Section III we propose some heuristics in order to simplify the problem and provide sound solutions from a biological point of view. Validation of the results and timings are presented in Section IV. Finally, a summary of findings and directions for future research are discussed.

## II. THEORY

### A. Preliminaries

The problem under study concerns the processing of strings of symbols belonging to the DNA, however the solutions proposed could be applied to a generic finite alphabet of symbols. Without loss of generality we will use a genomic alphabet Σ_4_ composed by the set of the four symbols {*A, C, G, T*}. The alphabet used in common Fasta files can include other symbols. For the aim of this paper k-mers lower case letters will be considered as their corresponding upper case symbols, while k-mers containing any other symbol will be discarded, since they mark areas of inferior quality.

#### Definition 1

(extended alphabet). The genomic alphabet Σ_4_ can be extended with a symbol *N* that stands for any symbol not in *{A, C, G, T}*. Such alphabet will be called 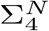.

#### Definition 2

(complementary symbols). Each symbol of the genomic alphabet Σ_4_ has a complementary symbol which is univocally identified by the function *compl*: Σ_4_ → Σ_4_, defined as *compl*(*A*) = *T*, *compl*(*C*) = *G*, *compl*(*G*) = *C*, *compl*(*T*) = *A*.

#### Definition 3

(word, k-mer). Let 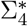 and 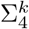 be, respectively, the set of all the strings of finite length and of length *k* over the genomic alphabet. A word is a finite ordered sequence of symbols 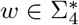. The length of a word is denoted by *|w|*. A word of length *k*, 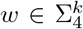, is called *k-mer*. The *j*th symbol of a k-mer is referred to as *σ_j_*, for *j* = 1*, …, k*.

#### Definition 4

(inverse-complementary word). For any word *w* = *σ*_1_*σ*_2_ … σ*_k_*, we define its inverse-complementary word 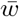 as

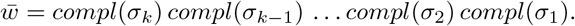

#### Definition 5

(reference Genome, set of k-mers of a reference Genome). A word that is as long as the genetic code of an entire organism is called “reference Genome” and is denoted by 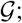 the set of its k-mers is indicated by 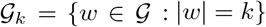, clearly 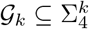.

#### Definition 6

(Hamming distance). The Hamming distance between two k-mers *w* and *w*′ is defined as *H*: 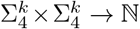

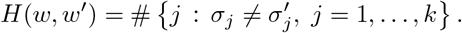

#### Definition 7

(distance of a word from a reference Genome). Given a reference Genome 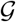 and its complete set of k-mers, 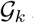, we define the distance of a word *w* from 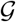 as

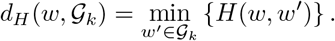

### B. The problem

A free text definition of the decision problem discussed in this paper could be “is there any string of length *k* (k-mer), composed of DNA symbols (*A, C, G, T*), that differs by at least *n* symbols from the complete set of k-mers of a reference genome?” This decision problem can be reformulated as an optimization problem by requiring that *n* is maximal.

Also the inverse-complementary words should be evaluated in the same genome, however in the following we shall ignore this latter hypothesis, since this can be easily taken into consideration without increasing the asymptotic complexity of the problem.

**Problem 8** (MAX-KMER problem, MAX-KMER decision problem). The MAX-KMER problem consists in identifying the maximally distant k-mer 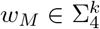, such that

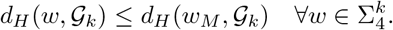

The corresponding decision problem is stated as follows.

Given *δ* ∈ ℕ and a word 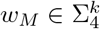

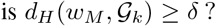

The MAX-KMER problem is 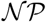-Hard; this can be easily proved noting that once we map genomic strings into binary strings we obtain an instance of the dual version of the Hamming Center Problem (HCP), which is 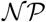-Hard [1].

#### Definition 9

(Hamming Center problem (HCP), Hamming Center decision problem (HCDP)). Let 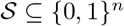 be a finite set of binary strings of length *n*. The HCP consists in finding a binary string *β*_*m*_ ∈ {0,1}^*n*^ such that

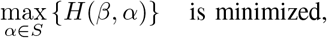

i.e. 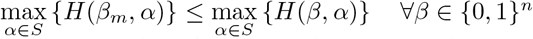.

The corresponding decision problem is stated as follows.

Given *δ* ∈ ℕ and a binary string *β*_*m*_ ∈ {0,1}^*n*^

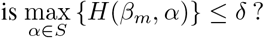

#### Theorem 10. The MAX-KMER problem is 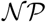-Hard

*Proof:* We prove the 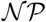-hardness of the MAX-KMER using a polynomial reduction from the Hamming Center problem. We define a correspondence between a k-mer *w* = *σ*_1_ *… σ_k_* and a binary string by means of a bijective map *inBinary*(*w*): 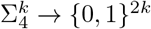
where *b*: Σ_4_ → {0, 1}^2^ is a map defined as *b*(*A*) = 00, *b*(*C*) = 01, *b*(*G*) = 10 and *b*(*T*) = 11.

The inverse map *fromBinary*: 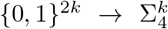 it is defined as

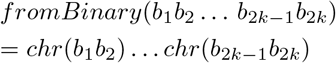

where *chr*:= *b*^−1^: {0, 1}^2^ → Σ_4_, *chr*(00) = *A*, *chr*(01) = *C*, *chr*(10) = *G* and *chr*(11) = *T*.

To solve the HCP problem using an algorithm for the MAXKMER problem we express the HCP, defined on an input set of binary strings 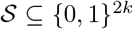, in its dual form and then use the map *fromBinary* to reformulate the problem as a MAX-KMER over the genomic alphabet. The HCP is equivalent to its dual form in which we maximize the minimum number of matches. The problem consists in finding a binary string *β*_*m*_ ∈ {0,1}^*n*^ such that 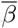 is the binary complement of *β*, i.e. is a string of the same length of *β* in which each binary symbol is replaced by its complement (1 by 0 and vice versa). Let *β*_*m*_ be a solution of the HCP that solves also the dual problem. By taking *fromBinary*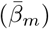 we obtain a solution of the MAX-KMER problem.

Conversely, let *w*_*M*_ be a solution of the MAX-KMER problem, by taking *inBinary*(*w*_*M*_) we obtain *β*_*M*_, and 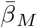 is the solution of an instance of the HCP.

**Proposition 11.** *The MAX-KMER decision problem is 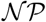-Complete*.

*Proof:* to obtain a polynomial certificate for a MAX-KMER decision problem solution *〈*w*_*M*_, δ〉* it is sufficient to search the k-mer *w*_*M*_ in the complete set of k-mers 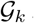 of the reference genome 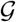, in 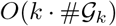, and report the minimum Hamming distance *m*; if *m* ≤ *δ* accept the solution, otherwise reject it.

## III. IMPLEMENTATION

### A. Algorithm

The algorithm for searching the current candidate against the reference genome is linear in the size of the genome; the pseudo-code is shown in Algorithm 1. It takes as inputs the reference genome and the current candidate sequence *curr*_*seq*. For each offset in the reference genome it computes in parallel, using GPU cores or CPU threads, the arrays of the differences between the current direct and inverse complementary sequences, named **dd** and **di** respectively; the elements of the arrays are filled according to internal scheduling of the threads, whose index is indicated by *thr*_*id*. The inverse complement of the current sequence, *seq*_*i*, is determined at line L.4. Then, two parallel reductions, implemented again using GPU cores or CPU threads, are applied to the arrays of the differences in order to obtain the global minima *md* and *mi*. The global minima are finally inserted into a max priority queue that performs a Pareto optimal comparison of both criteria. Candidate sequences can be explored in several ways, however it is better to generate them by enumeration, so we obtain two advantages: First, by establishing a total order among k-mers we do not need to keep track of the candidates already processed and so we can build an anytime algorithm that can be interrupted and restarted. Second, also the pairs of a sequence and its inverse complementary are ordered, therefore we can immediately tell, by establishing a conventional order, whether the current candidate is a new code or the inverse complementary of an already processed code; lines L.5-L.8 of Algorithm 1 exploit this fact by skipping pairs that have already computed.

#### Algorithm 1

parallel algorithm for searching a k-mer against a reference genome.

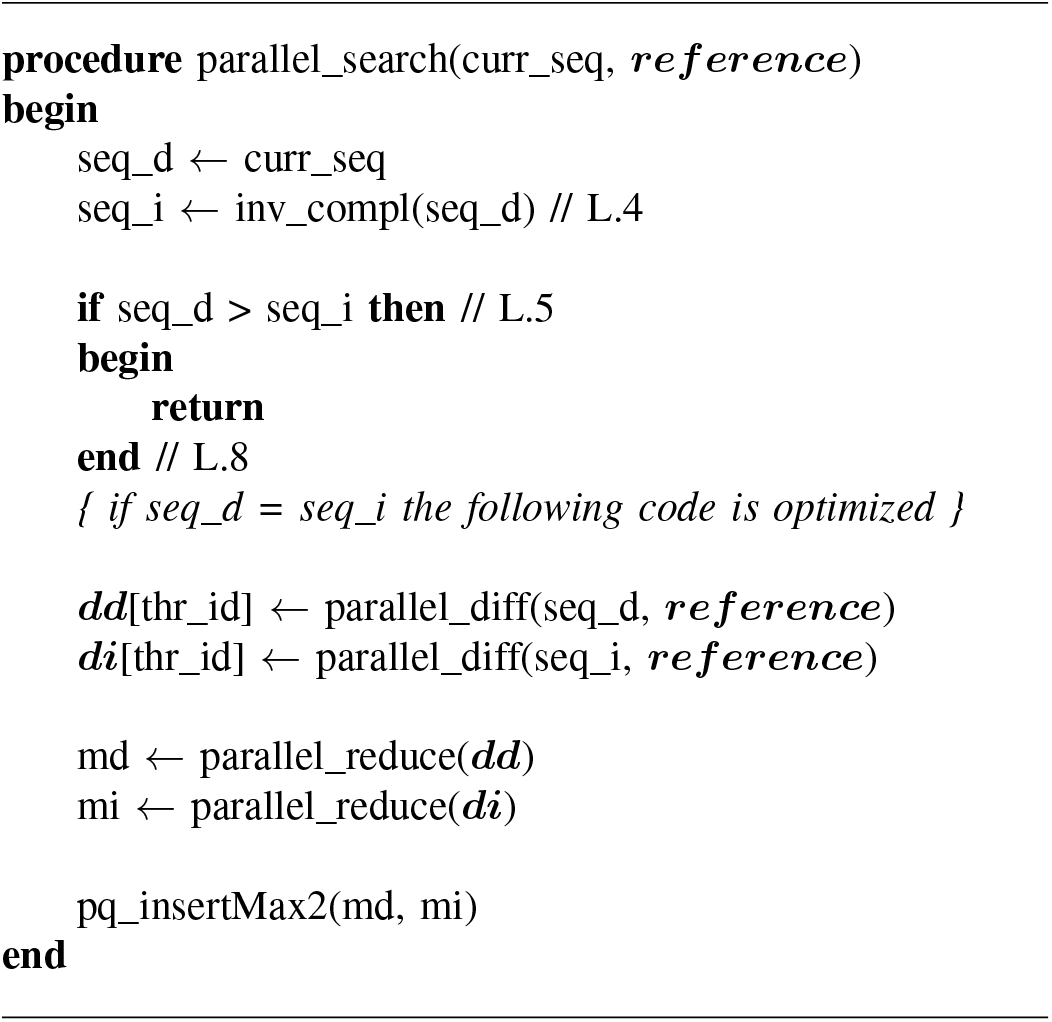

In the case we want to fix the number of occurrences of each symbol, another type of exhaustive generation of k-mers is represented by the set of the permutations of a initial sequence. Over permutations it is indeed possible to establish a total order by relating them to a factorial number system, i.e. a mixed radix numeral system adapted for numbering permutations [2]. This could have a meaning from a biological point of view since, for example, it is possible to fix *a priori* the percentage of *C* and *G* symbols (see section III-C2).

#### Algorithm 2

algorithm for determining the number of different symbols between two 64-bit binary numbers.

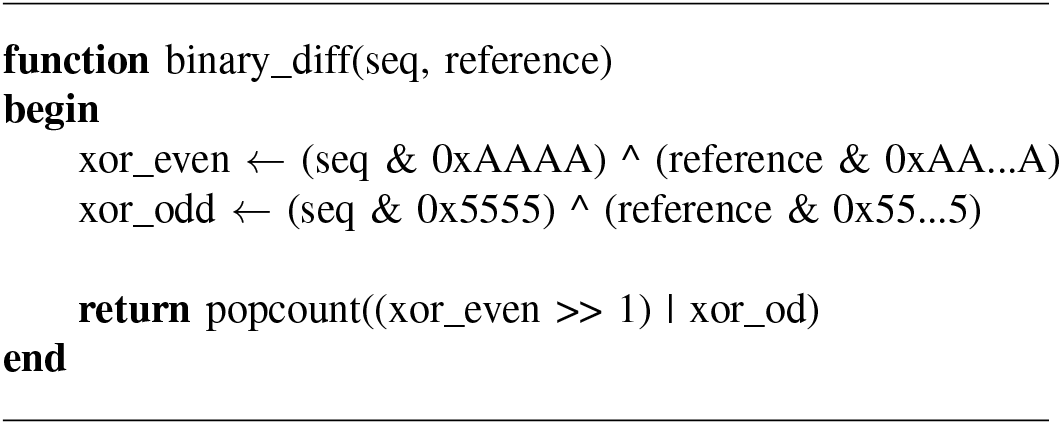

### B. Representation of the genome words

Genomes are huge sets of data, for instance the human genome has about 3 × 10^9^ base pairs. For this reason we will operate on bits in the domain 0, 1 to represent the 4 main symbols in Σ_4_. If we consider that the ASCII standard uses 8 bits to represent a single character while 4 symbols can be stored with *log*_2_(4) = 2 bits of information, it is easy to figure a gain of 75%; in the real case the reference alphabet is 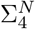, therefore we have to use 4 bits and the gain is reduced to 50%. We cannot use simply 3 bits to store the additional symbol because there would be slack bits in the 64-bit registers and the symmetry and efficiency of the algorithms would suffer. We consider the lexicographical order of the symbols: A, C, G, T and we assign to each symbol a progressive base 2 number to represent it, therefore we use the numbers 00_2_, 01_2_, 10 and 11_2_ respectively. In practice, we are using half a byte, that is a nibble, per symbol; an example for 4-mers is reported in Table I.

**Table I.** POSSIBLE REPRESENTATIONS OF 4-MERS.

| String | Binary | Base 4 | NibbleHex |
| --- | --- | --- | --- |
| AAAA | 00 00 00 00 | 0000 | 00 |
| AAAC | 00 00 00 01 | 0001 | 01 |
| AAAG | 00 00 00 10 | 0002 | 02 |
| AAAT | 00 00 00 11 | 0003 | 03 |
| ... | ... | ... | ... |
| AATA | 00 00 11 00 | 0030 | 0C |
| ... | ... | ... | ... |
| ATAA | 00 11 00 00 | 0300 | 30 |
| ... | ... | ... | ... |
| TAAA | 11 00 00 00 | 3000 | C0 |
| ... | ... | ... | ... |
| TTTT | 11 11 11 11 | 3333 | FF |

The binary representation allows for the use of bitwise operations that are directly mapped on atomic processor operators at hardware level, being performed on 64-bit registries, as illustrated in Algorithm 2.

The operators &,|, ^ and *>>* stand for bitwise “and”, “or”, “xor” and “right shift” respectively. The only non atomic operation is the function *popcount* for counting the number of ones in the sequence, however it is one of the SSE v4.2 intrinsic instructions provided by modern CPUs and so it is very fast. The search algorithm has also been written for Nvidia GPUs exploiting their massive number of computational cores and their complex memory hierarchy: The reference genome is stored in global memory, while the candidate sequence is kept in the constant memory; the latter is optimized at hardware level for efficient broadcasting towards all cores. The reduction operations are implemented exploiting the shared memory and require a logarithmic number of steps [RIF CUDA].

### C. Heuristics and good biological solutions

Since the problem is 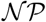-hard, under the hypothesis that DNA fragments contain information a heuristic criterion is to increase the “entropy” of the sequences. Moreover, even the optimal solution could not be satisfactory from a biological point of view: Primers have indeed features that characterize their “fitness”, and these features can be exploited to reduce the candidate k-mers.

#### 1) Abstract heuristics

A simple heuristics consists in promoting k-mers with a higher heterogeneity of symbols. Real genomes are very complex, therefore we have to choose relatively long values for *k* in order to have a reasonable number of missing combinations. To have an idea about which values, we built a hash table containing all k-mers of three reference genomes: A small bacterial genome (*Mycobac-terium tubercolosis*, 4MB), a small plant genome (*Arabidopsis thaliana*, 120MB) and the human genome (3GB); in Table II we report three measures, redundancy, relative redundancy and coverage, defined as in the following.

**Table II.** SOME STATISTICS.

|  |  | M. tuberculosis | A. mediterranei | A. thaliana | Pyrus sp. | X. tropicalis | H. sapiens |
| --- | --- | --- | --- | --- | --- | --- | --- |
| **Word size 10**<br>(tot. 1,048,576) | **Singleton codes** | 195,035 | 163,072 | 1,872 | 0 | 0 | 0 |
|  | **Total codes** | 742,505 | 657,916 | 1,047,819 | 1,048,576 | 1,048,576 | 1,048,576 |
|  | **10-mers** | 4,411,522 | 10,236,705 | 118,960,596 | 501,311,979 | 856,997,629 | 2,864,785,127 |
|  | **Maximum freq.** | 437 | 1,108 | 54,523 | 677,362 | 27,494 | 2,975,193 |
|  | **Coverage** | 70.81% | 62.74% | 99.93% | 100.00% | 100.00% | 100.00% |
|  | **Rel. redundancy** | 81.40% | 84.45% | 99.82% | 100.00% | 37.26% | 100.00% |
|  | **Redundancy** | 83.17% | 93.57% | 99.12% | 99.79% | 99.88% | 99.96% |
| **Word size 15**<br>(1,073,741,824) | **Singleton codes** | 4,073,268 | 7,702,781 | 71,513,541 | 131,846,345 | 187,112,889 | 177,353,773 |
|  | **Total codes** | 4,211,727 | 8,755,427 | 87,734,907 | 212,247,286 | 370,992,577 | 546,557,336 |
|  | **15-mers** | 4,411,517 | 10,236,700 | 118,960,591 | 501,311,974 | 856,997,624 | 2,864,785,122 |
|  | **Maximum freq.** | 160 | 65 | 16,770 | 412,533 | 1,130 | 1,198,752 |
|  | **Coverage** | 0.39% | 0.82% | 8.17% | 19.77% | 34.55% | 50.90% |
|  | **Rel. redundancy** | 99.62% | 99.28% | 99.05% | 87.72% | 100.00% | 83.48% |
|  | **Redundancy** | 4.53% | 14.47% | 26.25% | 57.66% | 56.71% | 80.92% |
| **Word size 20**<br>(1,099,511,627,776) | **Singleton codes** | 4,298,788 | 10,085,966 | 106,272,968 | 291,987,116 | 742,478,611 | 2,157,756,102 |
|  | **Total codes** | 4,339,979 | 10,153,705 | 110,093,809 | 340,528,917 | 784,031,866 | 2,267,445,153 |
|  | **20-mers** | 4,411,512 | 10,236,695 | 118,960,586 | 501,311,969 | 856,997,619 | 2,864,785,117 |
|  | **Maximum freq.** | 52 | 53 | 10,147 | 311,195 | 872 | 451,296 |
|  | **Coverage** | 0.00% | 0.00% | 0.01% | 0.03% | 0.07% | 0.21% |
|  | **Rel. redundancy** | 100.00% | 100.00% | 100.00% | 99.97% | 100.00% | 99.80% |
|  | **Redundancy** | 1.62% | 0.81% | 7.45% | 32.07% | 8.51% | 20.85% |

##### Definition 12

(genome multiset). We define the multiset of a genome 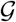 as

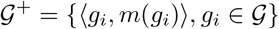

where *m*: Σ^*k*^ → *N* is the usual generalized multiplicity function [RIF?].

##### Definition 13

(redundancy). the redundancy of a set of k-mers 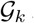 is

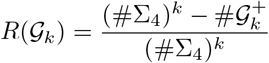

##### Definition 14

(relative redundancy). the redundancy of a set of k-mers 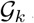 is

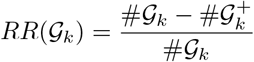

##### Definition 15

(coverage). the coverage of a set of k-mers 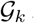 over an alphabet Σ_4_ is

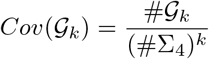

It is clear from Table II that 15 is a reasonable number for interesting genomes such as the human genome.

In any case, since the majority of Genome words should have statistically a normal distribution of symbols, the heuristic would measure in some way the “heterogeneity” of their composition and to prioritize candidate words with a higher (or smaller, but, again, this would be of little interest from a biological point of view) heterogeneity. In Figure III.1 we show the distribution of one of the complexity measures discussed below for the Human genome.

**Figure III.1.**
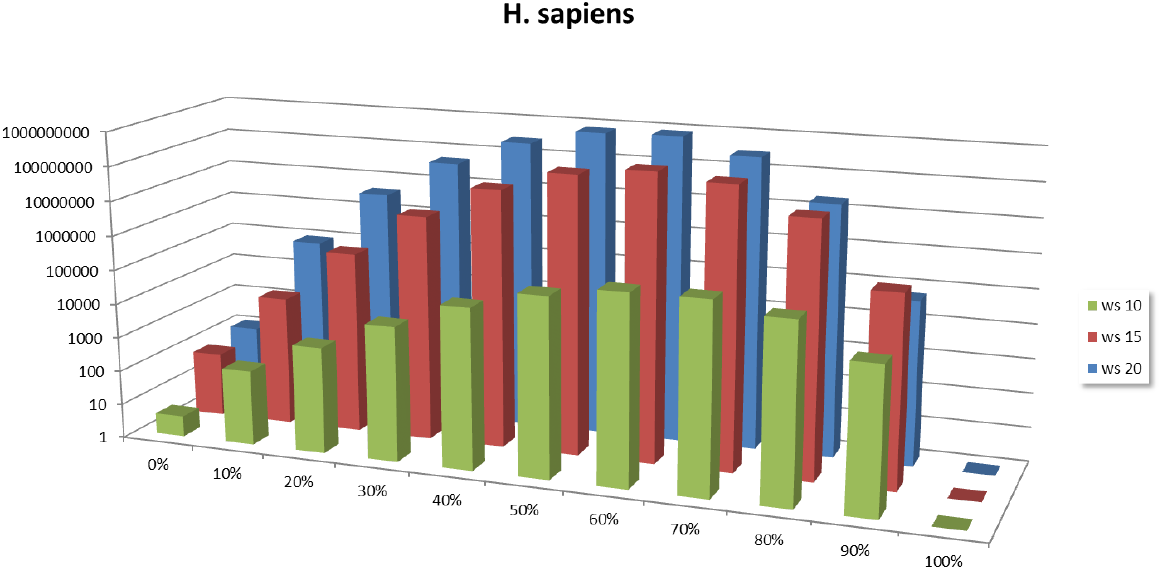
distributions of the positional complexity in the human genome for 10-mers, 15-mers and 20-mers.

The classical definition of heterogeneity is in terms of information entropy [3] defined as

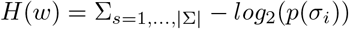

where *p*(*σ*_*i*_) is the frequency of the symbol *σ*_*i*_. A more “refined” idea comes from Wootton [4], [5] and allows evaluating the local compositional complexity for a word of length *k*:

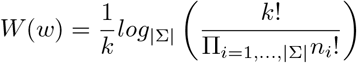

where *n*_*i*_ is the number of occurrences of the *i*^*th*^ symbol in *w*.

The previous formulas for *H*(*w*) and *W* (*w*) are not aware of the positions of the symbols in the word, only of their frequencies. For this reason, we developed a “positional complexity” that takes into account all the sequences of symbols in the word occurring every 1*, …, k* 1 symbols; the positional complexity seems more effective in describing the complexity of a given set of words; in Figure III.2 we can observe how its pattern is more variate with respect to those of the classical entropies and the Wootton’s one. However, although satisfactory from a theoretical point of view the positional complexity, as well as its relatives, is too slow in practical contexts, being *O*(*k*^3^). A faster alternative, that provides similar gains, relies on the comparison of just 4 symbols, for a total of 6 combinations, as shown in Algorithm 3. It is denoted as “Diff4” in Figure III.2.

**Figure III.2.**
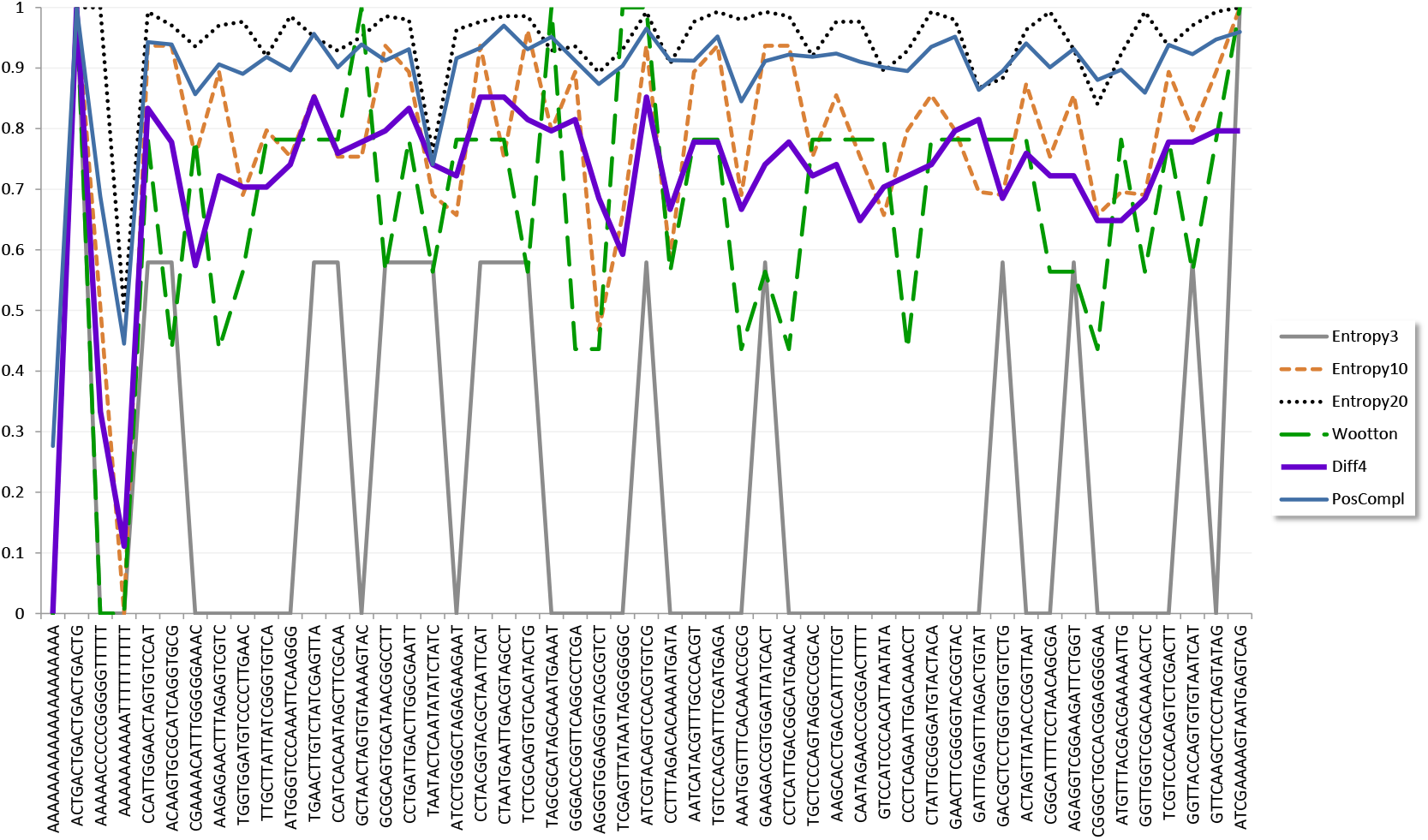
several definitions of entropy.

#### 2) Biological selection criteria

To reduce the search space we can rely on some desired properties of k-mers when contextualized in a biological environment. An easy criterion is the detection of series of consecutive identical symbols: It is possible to set a threshold and discard k-mers containing too long series that would cause alignment shifts. Also k-mers ending in “AA”, “AT”, “TA” and “TT” can be discarded, as well as those containing more than 3 symbols in *C, G* in the last 5 positions.

If a k-mer contains genomic palindromes it can form hairpins or self-dimers because of the presence of self-complementary regions within its sequence or between couples of identical sequences, therefore they should be avoided.

##### Algorithm 3

simplified version of the positional complexity.

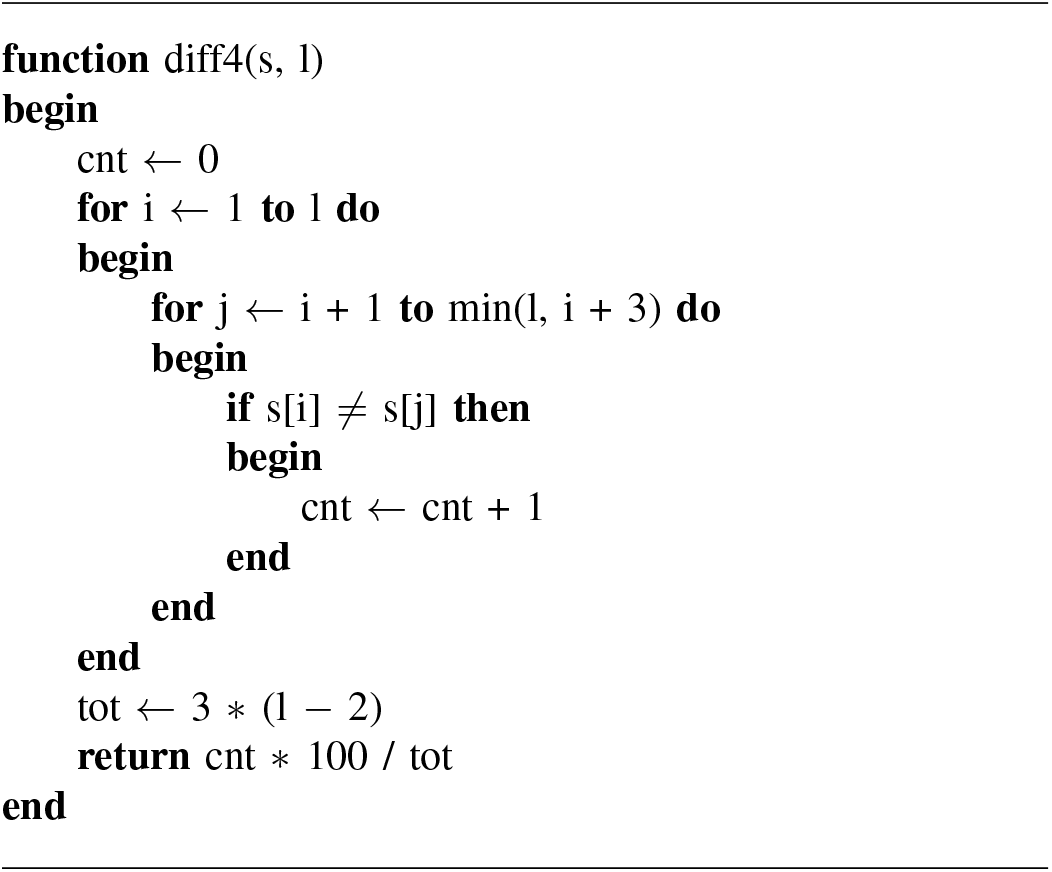

##### Definition 16

(genomic palindrome). A word *w* = *σ*_1_ *… σ_k_* is a (genomic) palindrome if it satisfies the function *gPalin*: Σ^*k*^ → {FALSE, TRUE},

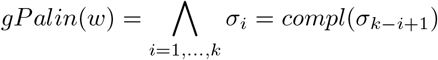

We can use a “longest common subsequence” algorithm (LCS) and compute all possible alignments between a sequence and itself. If the length of the common subsequence exceeds a given threshold the sequence is discarded. An additional check is done on the ends of the sequences: if the last position in the alignment is a match, and four out of the last five positions are matches, the sequence is discarded.

A balanced GC content is essential for a primer to be functional. For this reason it is possible to limit the GC content of a k-mer, usually between 40% and 60%.

##### Definition 17

(GC content). The GC content of a word *w* = *σ*_1_ *… σ_k_* is given by the function *gcContent*: Σ^*k*^ → *N*

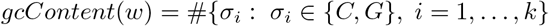

A final filter is computed on the so-called “melting temperature”. The melting temperature of a candidate k-mer is calculated with the Nearest Neighbor method and the SantaLucia table with DNA/DNA thermodynamic parameters, [SantaLucia, et al., 1996]:

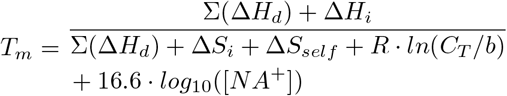

. ∆*H*_*d*_ and ∆*S*_*d*_, sums of enthalpy and entropy respectively, are calculated for all internal nearest-neighbor doublets; ∆*S*_*self*_ is the entropic penalty for self-complementary sequences. ∆*H*_*i*_ and ∆*S*_*i*_ are the sums of initiation enthalpies and entropies, *R* is the gas constant and *C*_*T*_ is the molar concentration. *b* is a constant equal to 4 for non-self-complementary sequences and 1 for duplexes of self-complementary strands or for duplexes when one of the strands is in significant excess.

All these combined filters allow for a drastic reduction of plausible candidates, as illustrated in Table III for permutations. Note that the total number of permutations is complete, being equal to their theoretical number given by the multinomial coefficient

**Table III.** FILTERED SEQUENCES FOR SEVERAL COMPOSITIONS OF SYMBOLS

| Initial permutation | passed | total | % | theoretical | ms | μs per seq |
| --- | --- | --- | --- | --- | --- | --- |
| 5:5:5:5 | 314,531,127 | 11,732,745,024 | 2.68% | 11,732,745,024 | 26,289,100 | 2.24 |
| 5:4:5:4 | 37,598,833 | 771,891,120 | 4.87% | 771,891,120 | 1,600,000 | 2.079 |
| 4:4:4:4 | 3,550,455 | 63,063,000 | 5.63% | 63,063,000 | 151,134 | 2.40 |
| 3:3:3:3 | 208,271 | 369,600 | 56.35% | 369,600 | 520.512 | 1.41 |
| 2:2:2:2 | 1,695 | 2,520 | 67.26% | 2,520 | 3.11 | 1.84 |

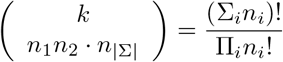

where *n*_*i*_ is the number of occurrences of the *i*^*th*^ symbol, written as *n*_*A*_: *n*_*C*_: *n*_*G*_: *n*_*T*_ in the table.

## IV. RESULTS

### A. Validation

The algorithm has been implemented in a tool named keeSeek (https://bitbucket.org/mfalda/cuda_keeseek/) implemented in CUDA. We ensure the correctness of the proposed tool by validating its results against glsearch (version 36.3.5b) [6], a global-local aligner part of the Fasta3 package and based on the Needleman and Wunsch algorithm [7]. Current software for sequence alignments is designed to search for the highest similarity between two sequences possibly by inserting gaps between symbols, while keeSeek is designed to search for the highest dissimilarity among continuous sequences. Nonetheless, glsearch can be used to confirm keeSeek results if candidate k-mers do not present discontinuities. The glsearch command line arguments are:

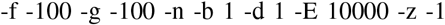

. To make the results from keeSeek and glsearch comparable, we try to discourage the presence of gaps by heavily penalize gap extensions (-g parameter). Even if performance was not the target of these comparisons, we reported also glsearch times in the next tables (Table IV and Table V).

**Table IV.** EXPECTED TIMES TO PRODUCE THE FIRST 128 FILTERED SEQUENCES USING ENUMERATION ON DIFFERENT GENOMES.

| Reference genome | Genome size | length | server1 | server2 | desktop1 | notebook1 |
| --- | --- | --- | --- | --- | --- | --- |
| *Mycobacterium tuberculosis* | 4.4 MB | 20 | 0m2.7s | 0m1.1s | 0m0.408s | 0m23.81s |
| *Amycolatopsis mediterranei* | 10.2 MB | 20 | 0m58.3s | 0m2.0s | 0m1.036s | 0m54.08s |
| *Arabidopsis thaliana* | 120 MB | 20 | 11m27.2s | 0m12.9s | 0m9.937s | 7m0.889s |
| *Pyrus sp.* | 500 MB | 20 | 48m25.42s | 0m50.59s | 0m41.847s | NOT POSSIBLE |
| *Xenopus tropicalis* | 1.5 GB | 20 | 2h19m14s | 2m2.87s | 2m2.168s | NOT POSSIBLE |
| *Homo sapiens* | 3 GB | 20 | 4h51m22s | 6m12.8s | NOT POSSIBLE | NOT POSSIBLE |

**Table V.** EXPECTED TIMES TO PRODUCE THE FIRST 128 FILTERED SEQUENCES USING PERMUTATIONS ON DIFFERENT GENOMES.

| Reference genome | Genome size | composition | server1 | server2 | desktop1 | notebook1 |
| --- | --- | --- | --- | --- | --- | --- |
| *Mycobacterium tuberculosis* | 4.4 MB | 5:5:5:5 | 0m2.5s | 0m1.1s | 0m0.612s | 0m23.28s |
| *Amycolatopsis mediterranei* | 10.2 MB | 5:5:5:5 | 0m58.1s | 0m1.6s | 0m0.868s | 0m53.99s |
| *Arabidopsis thaliana* | 120 MB | 5:5:5:5 | 11m16.0s | 0m13.4s | 0m9.553s | 7m2.203s |
| *Pyrus sp.* | 500 MB | 5:5:5:5 | 48m19.79s | 0m52.83s | 0m41.843s | NOT POSSIBLE |
| *Xenopus tropicalis* | 1.5 GB | 5:5:5:5 | 2h19m16s | 2m2.84s | 2m1.396s | NOT POSSIBLE |
| *Homo sapiens* | 3 GB | 5:5:5:5 | 4h52m02s | 6m00.4s | NOT POSSIBLE | NOT POSSIBLE |

### B. Performance

To measure the performance in terms of speed, we consider the three different reference genomes plus other three: A medium sized bacterium (*Amycolatopsis mediterranei*, 10MB), a small plant (*Pyrus sp.*, 500MB) and a vertebrate (*Xenophus tropicalis*, 1.5GB).

Four systems have been tested:

- **server1:** Intel Xeon E5540 quad core processors, 2.53GHz, 32GB RAM;
- **server2:** AMD Opteron 6128 quad core processors, 2.6GHz, 64GB RAM with a GPU Nvidia Fermi M2050, 6GB global memory;
- **desktop1:** Intel Q6600, 2.40 GHz, 3GB RAM with a GPU Nvidia GeForce GT 640 GDDR5, 2GB global memory;
- **notebook1:** Intel i7 M260, 2.67 GHz, 4GB RAM with a GPU Nvidia GeForce 310M, 0.5GB global memory.

Tables IV and V show the times required to search 128 20-mers in several genomes using candidates generated by enumeration and by permutation respectively. Additional time is required to load reference genomes (order of minutes for the bigger genomes).

## V. CONCLUSIONS

Finding non-existent words in genomes is an important problem of Computational Biology. We have proposed a tool for finding a set of sequences whose minimal Hamming distance from a reference genome is guaranteed; it also provides information about the location of the best solutions in the genome.This problem is not easy and we have formally proved this fact. We have also implemented a software in C++ exploiting the massive parallelism of modern Graphical Processing Units.

The problem under study can be solved using distributed algorithms and we intend to write a empowered version using MPI technology. Another possible enhancement is to assign a weight to each mismatch in order to rank solutions having the same distance from the reference genome.

